# AFsample3: Generating and selecting multiple conformational states with Alphafold3

**DOI:** 10.64898/2026.01.16.699904

**Authors:** Yogesh Kalakoti, Björn Wallner

**Affiliations:** Division of Bioinformatics, Department of Physics, Chemistry and Biology, Linköping University, 581 83 Linköping, Sweden; Science for Life Laboratory, Linköping University

**Keywords:** Sampling, Alphafold, Conformations, Ensembles, Diversity

## Abstract

Accurately capturing the conformational diversity of proteins is essential for understanding their mechanisms and regulation. However, current structure prediction approaches, including AlphaFold derivatives, are largely limited to modeling one dominant conformation. Improved sampling methods have expanded this to predicting two states. Here, we present AFsample3, an enhanced sampling framework built upon AlphaFold3 that substantially improves the generation and selection of diverse protein conformations. Across a benchmark of 238 non-redundant proteins with multiple experimentally determined states, AFsample3 significantly outperforms AlphaFold3 and its predecessor AFsample2 by improved predictions for 28% (67/239) of targets (ΔTM > 0.1) while degrading only 3% (8/239), and increases the number of high-quality alternate-state models (TM > 0.8) by 54% (from 54 to 83; p < 0.0001). AF-sample3 improves alternate-state accuracy for 28% of targets (ΔTM > 0.1) while degrading only 3%, and increases the number of targets with high-quality predictions (TM > 0.8) by over 50% compared to standard AlphaFold3. Ensemble diversity is also markedly enhanced (p < 0.0001), enabling accurate modeling of potential intermediate and or additional states. These results demonstrate that the improved sampling in AFsample3 can capture multiple native-like conformations, representing a significant advance in modeling of protein conformational landscapes.

## Introduction

Protein are critical macromolecules that serve as the building blocks of all life. They play crucial roles in almost all biological processes including metabolism, cellular communication, DNA replication, among others. In general, functional aspects of all proteins are highly dependent on their three-dimensional orientation. In solution, this 3D configuration is not static, instead, a protein system assumes different shapes (conformations) depending on factors including local environment, stage of a cellular process, cell-type, binding partners and others. While generative methods like Alphafold2 (AF2) (1) have been very effective in modeling protein structures, their default implementation lack the ability to generate conformational heterogeneity. To overcome this, sampling and perturbing input data representations of the Multiple sequence alignment (MSA) has proven to be very effective in inducing the generation of alternative conformations. These perturbations may be in the form of MSA sub-sampling (2), clustering (3) or masking (4, 5). Recently Al-phafold3 (AF3) was released as an update to the previous version that employs diffusion-based architecture, among other significant changes (6). AF3 can generate an all-atom model for a given query including nucleic acids, small molecules and ions, which is a substantial improvement to the AF2 inference system. Given this, it is important to probe the AF3 inference system for its ability to model multiple protein conformations.

Here, we present AFsample3, an alternate strategy based on the AF3 inference system for generating alternate protein states, and by extension, more diverse conformational ensembles. Like AFsample2 (5), it works by introducing noise in the inference system by randomly masking the input MSA representations at run time. The method was tested on 238 proteins having at least two distinct experimentally solved conformations (see Methods). We report significantly improved performance in both generating alternate states, and also generating an overall more diverse ensemble than baseline methods. For instance, AFsample3 was able to generate very high quality alternate state (TM-score>0.80) for 83 targets in the Cfold dataset, a substantial improvement over baseline (AF3vanilla: 54, AF2vanilla: 32), as well as existing methods (MSAsubsampling: 67, AFsample2: 63). Moreover, AFsample3 also reports a substantially increased level of conformational diversity (as quantified by fill-ratio) in the generated ensembles when compared to other methods. AFsample3 also improves the reference-free state selection system using a distance-scoring (DiSco) protocol, which, in principle, can identify an arbitrary number of conformations. Interestingly, the selection system identified potential novel conformations for multiple targets that satisfy known target-specific hypothesis’ in the literature. Taken together, AFsample3 provides an optimal system to generate and select protein conformations, without any additional network training on top of the AF2/AF3 architecture. Also, the developed system has practically no computational overhead than what is already required for the default AF3 inference system.

Given the architectural differences in AF3, these results and benchmarks provides important insight into whether perturbation strategies remain effective or require adaptation in the newer AF3 model. This study underlines key improvements to the AF3 inference system over AF2, particularly related to the robustness in performance on perturbed input data. Taken together, these results also serve as a reference on the predictive performance of both networks (AF2 and AF3) on a comprehensive dataset for the task of generating protein with multiple conformations.

## Methods

### Datasets

A recently curated dataset named *Cfold* (7) that compiled targets with alternate conformations in the Protein Data Bank (PDB) was utilized in this study. The dataset contains two states for 238 proteins that are not similar than TM>0.8, represented by the structures with the largest difference in TM-score, in the case the protein has more than two states. Figure 1 summarizes the overall distribution of reference structure similarities as well as sequence lengths. The conformational changes in TMscore ranges from around 0.3 to 0.8, denoting a wide range of conformational variability in the dataset. Also, the reference structures diverge more with increasing sequence length.

**Fig. 1.**
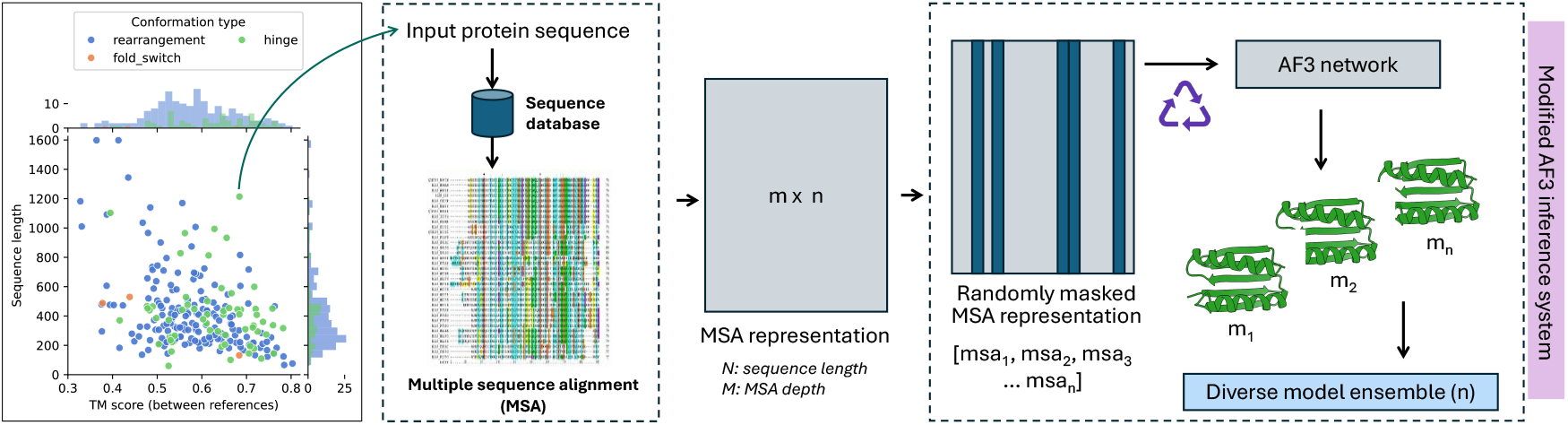
Overview of AFsample3 protocol. Given a dataset of proteins with varying degree of sequence length and TM-score (similarity) between known conformations, the AFsample3 protocol randomly masks multiple sequence alignment (MSA) in order to break evolutionary signals at each iteration of the inference run, making the system more susceptible to generating a diverse conformational ensemble.

### Structure generation

Standard and non-standard pipelines based on AF2 an AF3 were employed to generate models for the analysis. The default implementation of AF3 v3.0.1 was modified to have the functionality of MSA masking, as well as MSA subsampling, by restricting the *max_ali* parameter similar to what was done for subsampling in AF2 (2). The MSAs, which is the only input to all methods, were directly retrieved from Cfold Zenodo record (https://zenodo.org/records/10837082) and used as input for all predictions on all targets.

AFsample2 (wallnerlab.org/afsample2:v1.1) and AFsample3 (wallnerlab.org/afsample3:v1.0) was used to generate structures for the AF2, and AF3 networks, respectively, and for all the 24 different combinations of settings in Table 1. In total, 1000 models for each of the 24 settings and 238 targets in the Cfold datasets were generated, amounting to around 6 million models.

**Table 1.**
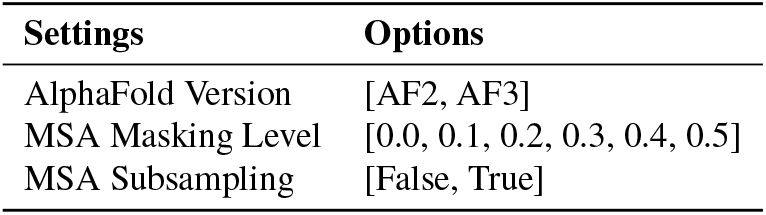
Set of parameters employed to study the effect of MSA perturbations in the different AlphaFold neural networks on the the ability to generate protein conformations.

| Settings | Options |
| --- | --- |
| AlphaFold Version | [AF2, AF3] |
| MSA Masking Level | [0.0, 0.1, 0.2, 0.3, 0.4, 0.5] |
| MSA Subsampling | [False, True] |

### End state analysis

TMalign (8) was used to calculate the similarity of the models to the available reference states to determine whether both conformations were present in the generated ensembles. The models most similar to the two end states, representing the best preferred and alternate conformation, respectively, were compared between each method, setting and ensemble. Ideally, a good method should be able to generate at least one good model (for example with TM-score>0.8) for both the conformations.

### Ensemble diversity

In order to investigate and compare conformational diversity of generated ensembles between methods, a visualization scheme called diversity plot was defined, where TM-scores of all models with the two reference states in a given ensemble are plotted against each other. This visualization provides an immediate sense of conformational preference for a given ensemble (Figure 6d). To quantify the level of diversity, our previously developed metric, *fill-ratio* was utilized (5). In short, the fill-ratio quantifies how much of the diagonal in the diversity plot is populated by models. It works by projecting all points in the diversity plot to a diagonal going between states and computing a weighted fraction of populated to unpopulated bins on the diagonal. The metric was updated so that it emphasizes the presence of both states by using a parabolic reweighting along the projected points, giving more weight to points closer to the end states.

### Sampling experiment

To determine how many models that needs to be generate to predict the alternate conformational states a sampling experiment was conducted. Given TM-scores, s0, and, s1, for two states for each target, the best possible score for the alternate state was extracted as the minimum of the maximum TM-score values, with a threshold set at 99% of this value. Random sampling with replacement was performed across sample sizes from 1 to 1,000 for multiple iterations, recording the smallest sample size yielding a similarity score (minTM) meeting the threshold. The experiment was bootstrapped for 100 iteration, estimating the minimal sampling requirements for conformational state recovery across methods.

### Statistical analysis

All pairwise comparisons were tested with the Wilcoxon rank sum test under the alternate hypothesis that one distribution is significantly higher than the other. SciPy v1.6.1 was employed to perform all statistical analysis.

## Results

Here we present AFsample3, the next iteration of the AFsample family of methods that are aimed at improving the conformational diversity in generated protein ensembles. AFsample3 is based on the Alphafold3 inference pipeline and utilize the same MSA sampling strategy, i.e. random column masking, as in AFsample2 (5). AFsample3 generates significantly better conformations for the end states, while at the same time improving the conformational diversity of the ensemble compared to all other methods in the benchmark. In the following sections we compile the results that highlight the improvements when compared to the AlphaFold baselines, AF2vanilla and AF3vanilla, as well as to existing state-of-the-art methods AF2conformations (2) (MSA subsampling) and AFsample2 (5). Additionally, we also compare ensemble diversity, sampling characteristics, and describe a method for reference-free state selection that can be applied to select more than two states.

### AFsample3 improves over AFsample2 and Alphafold3 for protein conformation prediction

All methods were evaluated on their ability to generate both conformational states of two-state protein systems. To eliminate variability due to differences in sampling, each method was used to generate 1,000 models per target. As previously mentioned, 12 different settings were tested across two network types for 238 targets in the Cfold dataset, resulting in approximately 6 million models (Table 1).

It is well established that AlphaFold-based methods tend to favor one conformational state in multi-state proteins, a trend also observed in the Cfold dataset (Figure 2b). Accordingly, for each target, the two conformational states were categorized as preferred or alternate based on preference of the default method. Figure 2 summarizes the ability of all methods to predict both states. As shown in Figure 2a, the default AF3 inference system (AF3vanilla) significantly outperforms AF2vanilla in generating both the preferred and alternate conformations. When comparing the quality of alternate states, AF3vanilla produced markedly better models (ΔTMscore ≥0.05) for 56 targets compared to AF2vanilla, while underperforming for only 23 targets. The remaining 159 targets (gray) showed marginal differences in quality of the generated alternate conformation (ΔTMscore *<* 0.05).

Interestingly, AF3vanilla performs only slightly worse than AFsample2 and generates better alternate conformations for 47 targets but performs worse for 52. However, AFsample3, using MSA masking, clearly outperforms all other methods in generating high-quality alternate states. Specifically, the improvement-to-deterioration ratios for alternate state prediction are 96:14, 67:8, and 72:20 when compared to AF2vanilla, AF3vanilla, and AFsample2, respectively.

**Fig. 2.**
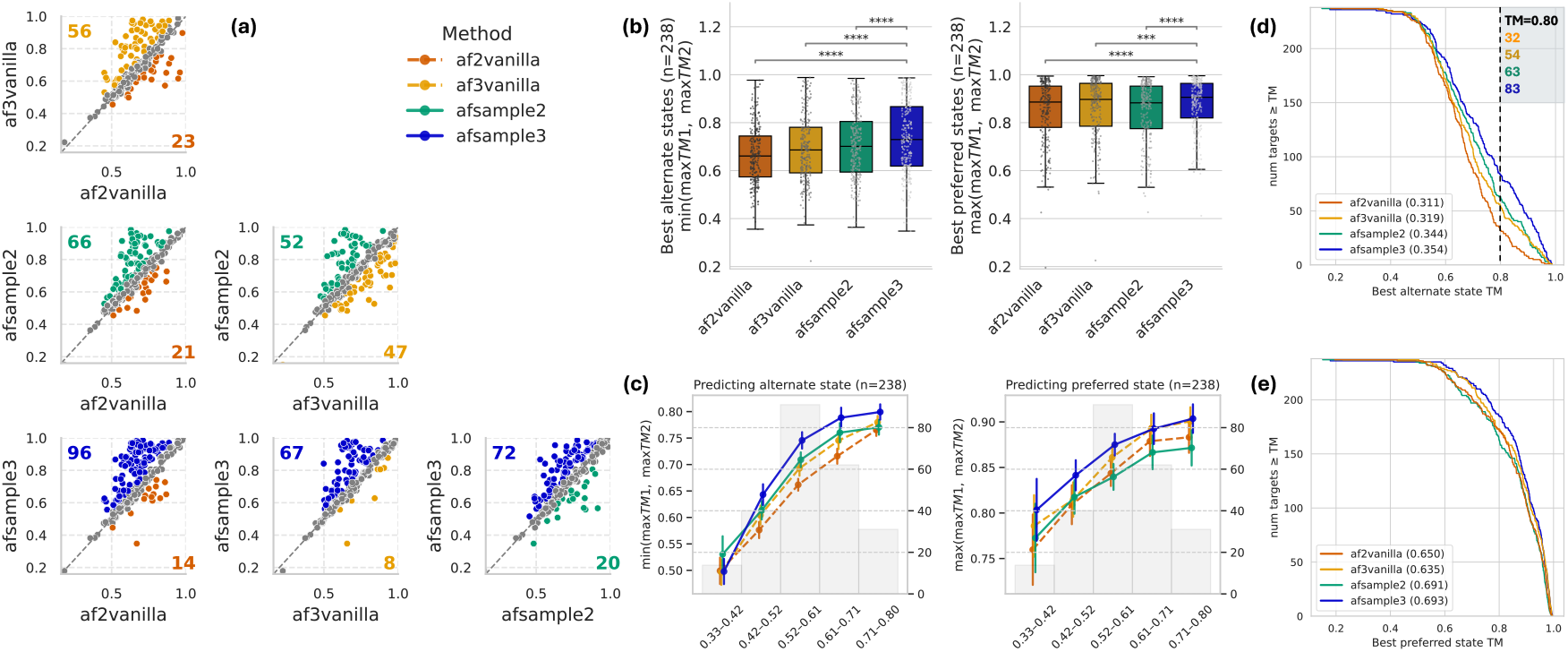
AFsample3 generates better end states for most targets in the Cfold dataset. (a) Comparing the quality of alternate state (TMScore) between four methods, namely, af2vanilla, af3vanilla, afsample2 and afsample3. It is clear that AFsample3 generates better end states for a majority of targets in the Cfold dataset. (b) The overall distribution of quality of both preferred and alternate states are summarized, and the improvements are clear and significant (Wilcoxon rank sum - ****: p<0.0001, ***:p<0.001). AFsample3 not only generates better alternate states (left), it also slightly improves the already good preferred state (right). This property is different from that of AFsample2 where the quality of the preferred state deteriorates marginally. (c) Analyzing the performance of targets with varying levels of conformational change. (d) Cumulative frequency distribution plots for generating the (d) alternate and (e) preferred conformations.

It is important to note that while AFsample2 often achieves improved alternate state modeling at the cost of slightly reduced quality for the preferred state (Figure 2b), AFsample3 does not exhibit this trade-off. Instead, it enhances the quality of both preferred and alternate conformations, highlighting its overall robustness.

Since the similarity of reference states varies considerably across targets, with TM-score in the [0.3,0.8] range (Figure 1), it was important to assess how performance depends on the reference state similarity. Figure 2c shows the differences in model quality between methods, grouped by bins of reference state similarity. The results indicate that AFsample3 performs consistently well across all target types, particularly for the majority of targets with reference states having a TM-score > 0.5. This is emphasized even more in the inverse cumulative distribution function shown in Figure 2d, where AF-sample3 generates very-high quality alternate state (TM>0.8) for 83 targets, much higher than AF3vanilla (54) and AFsample2 (63). It should be noted that AF3vanilla performs much better than AF2vanilla (Figure 2d), pointing to a combination of improved model architecture, training strategy, or a robust training set in Alphafold3.

### Effect of MSA randomization

AFsample3 improves sampling by randomly masking columns in the input MSA to the AlphaFold3 network. However, the optimal degree of randomization varies depending on the target or dataset. As with AFsample2, a range of randomization levels was tested for AF3. Figure 3a,b summarizes the average quality of the best generated states as a function of the randomization level for the Cfold dataset. In short, 20% and 40% randomization level for AF2 and AF3, respectively, yielded the best overall performance for generating both states. The 20% for AF-sample2 agrees with what was previously reported to be the optimal for a smaller set of 23 proteins (OC23) for AFsample2 (5). Unless otherwise stated, all reported results for AF-sample2 and AFsample3 use 20% and 40% randomization, respectively.

**Fig. 3.**
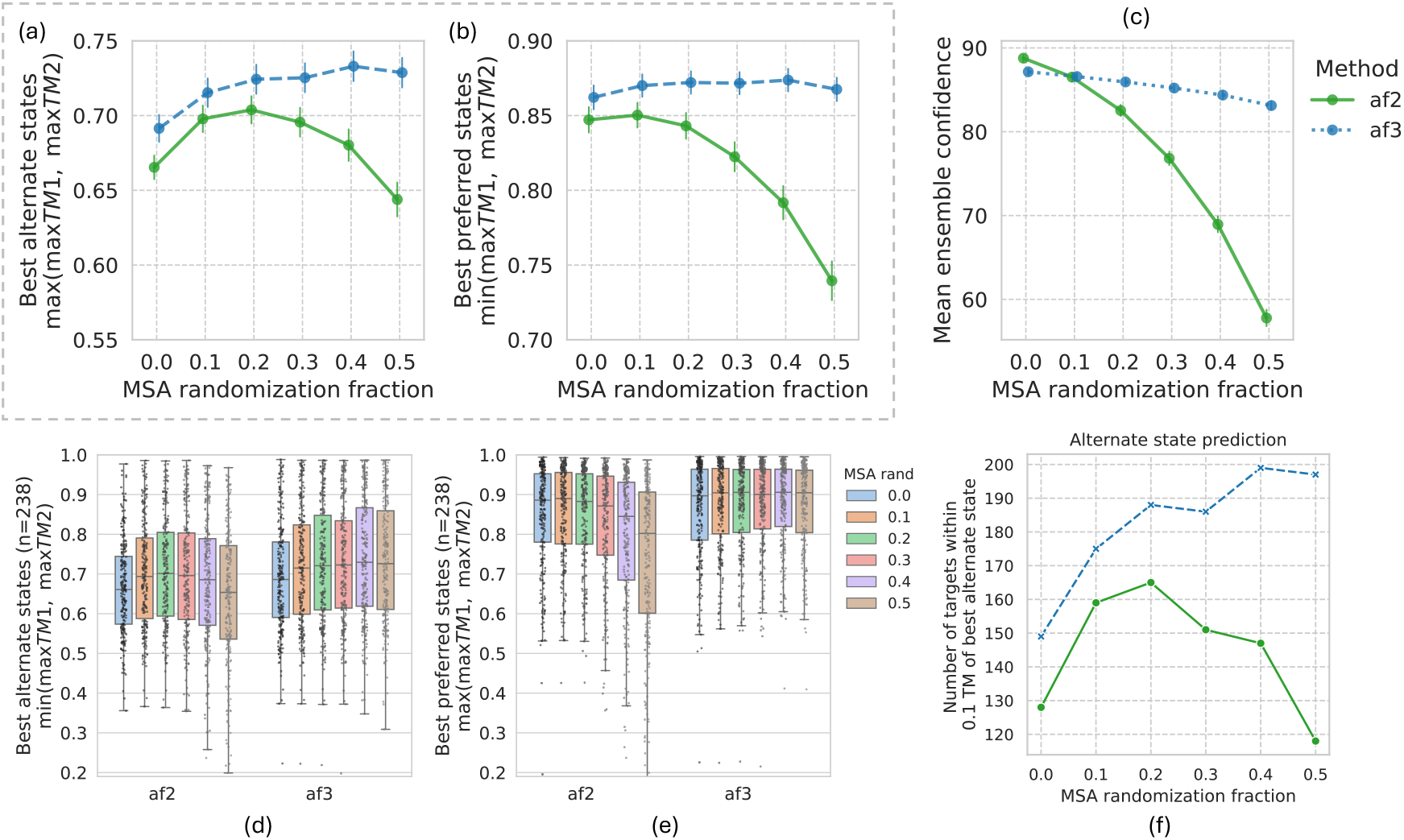
Effect of MSA perturbation on the ability of Alphafold2/3 to generate protein conformations for the preferred and alternate states in the Cfold dataset. Average quality for (a) the best preferred state and (b) best alternate state, (c) average model confidence, distribution of quality for (d) the preferred states and (e) alternate states at different randomization levels, (f) number of targets that is able to generate a model within 0.1 TM-score of the best alternate state generated from any setting for different randomization levels.

In addition, there are stark differences between AF2 and AF3 in response to MSA randomization, with AF2 being much more sensitive to MSA perturbation than AF3. For the preferred state, both AF2 and AF3 perform well with no MSA perturbation (average TM-score 0.85). In AF2, performance slightly improves at 10% randomization but then drops sharply (Figure 3b). By contrast, AF3 shows a small steady improvement up to around 40% randomization and does not exhibit the same performance decline as AF2.

Looking at the alternate states (Figure 3a), AF3 starts off significantly better than AF2 with no randomization and maintains its advantage across all randomization levels. A similar difference is also observed in the confidence scores (Figure 3c), the internal metric used to assess model quality, where the average confidence of the generated ensemble stays relatively stable and flat in AF3 when compared to AF2, which drops notably. Remarkably, AF3 maintains a high confidence level (>80) even when 50% of the MSA is randomized, this is might of course be related to the fact the AF3 models at this randomization levels are still quite good compared to the AF2 models. But it also underscore that AF3 is much more robust and tolerant towards MSA perturbations.

However, as observed in previous studies, the optimal level of MSA randomization is protein specific (5). While most models achieve their best performance at a particular randomization level, there are outliers for which both the preferred and alternate states are best generated at different levels of perturbation.

### Effect of sampling more models

It has been extensively shown that sampling plays a critical role in the ability of the methods to generate conformational states, and it remains a central component across all such approaches (6, 9, 10). In general, sampling more improves performance, although with diminishing returns as the number of samples increases. Thus, there is a trade-off between computational time and performance. To investigate this trade-off, the effect of varying the number of samples on performance was examined.

As discussed previously, both AF2 and AF3 have an inherent bias towards a preferred conformational state. Therefore, this analysis focused on how increased sampling affects the ability of each method to generate the alternate state. As shown in Figure 4, a sharp improvement in the quality of best alternate-state models is observed for both AFsample2 and AFsample3 up until 100 models, it still improves with more sampling but at a much slower rate. In general, AFsample3 consistently achieves a significantly higher mean TM-score (averaged across all targets) than AFsample2 at practically all sampling levels.

**Fig. 4.**
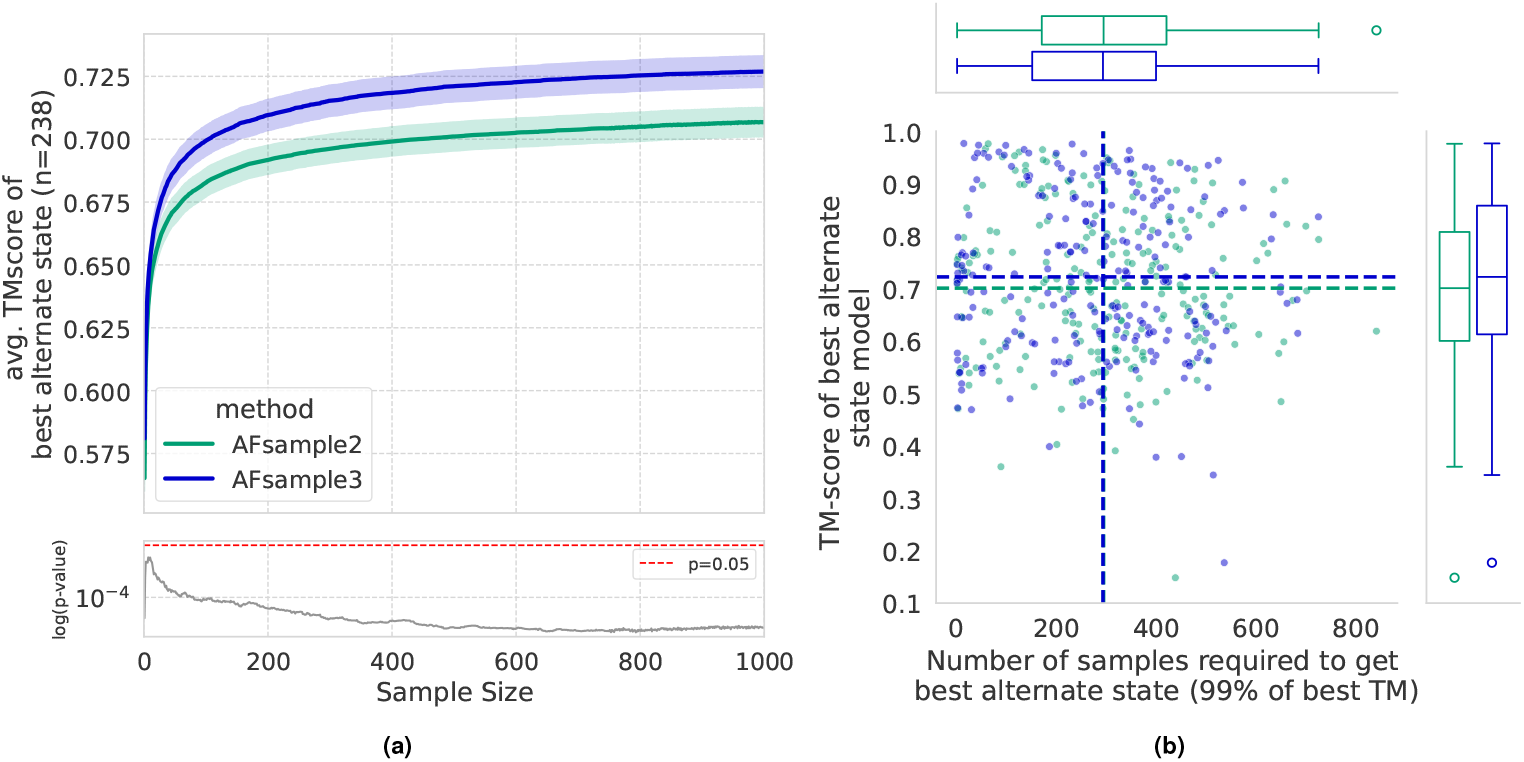
The importance of sampling in generating alternate states. (a) Quality of the best alternative states as a function of generated models. (b) Scatterplot depicting the amount of sampling required to generate the alternate state for each target in the Cfold dataset.

Again, similar to the case of MSA randomization level, the optimal sampling size is target-specific. As illustrated in Figure 4b, although the median sampling level for both AFsample2 and AFsample3 cluster around 300, there are clear outliers exist require significantly less or substantially more sampling to achieve optimal performance.

### Effect of MSA subsampling

MSA subsampling, where a shallower subset of the full MSA is used for inference, has previously shown strong performance with AF2 (2). In our earlier work, it consistently outperformed most alternative approaches and ranked second only to AFsample2 (5). Motivated by these results, and to explore the impact of reduced MSA depth on conformation generation in AF3, we implemented the MSA subsampling strategy in AF3 and compared the performance of the previous methods with and without MSA subsampling. Interestingly, the behavior of AF3 with subsampling is almost identical to AF2 with subsampling, significantly improving the generation of both the preferred and the alternate states (Figure 5). This indicates that a reduction in MSA depth influences conformational sampling in a comparable way across the two versions. Although AF3vanilla is slightly better than AF2vanilla, the improvement from subsampling is large: AF2 with subsampling outperforms AF3vanilla and performs very similarly to AF3 with subsampling for the best alternate state. Sub-sampling also improves AFsample2, while it does not improve AFsample3. In fact, subsampling with AFsample2 is on par with AFsample3, which highlights that combining different ways of MSA perturbations improves performance using the AF2 network. Two possible reasons for why AFsample3 does not benefit from subsampling could be that, the MSA track is smaller in the AF3 network and that the room for improvement is less since it is already the best performing method. Taken together, these findings show that sub-sampling substantially improves AF2vanilla, AF3vanilla, as well as AFsample2, but has no significant effect on AFsample3, which without subsampling remains the overall best-performing strategy for generating both the preferred and alternate states.

**Fig. 5.**
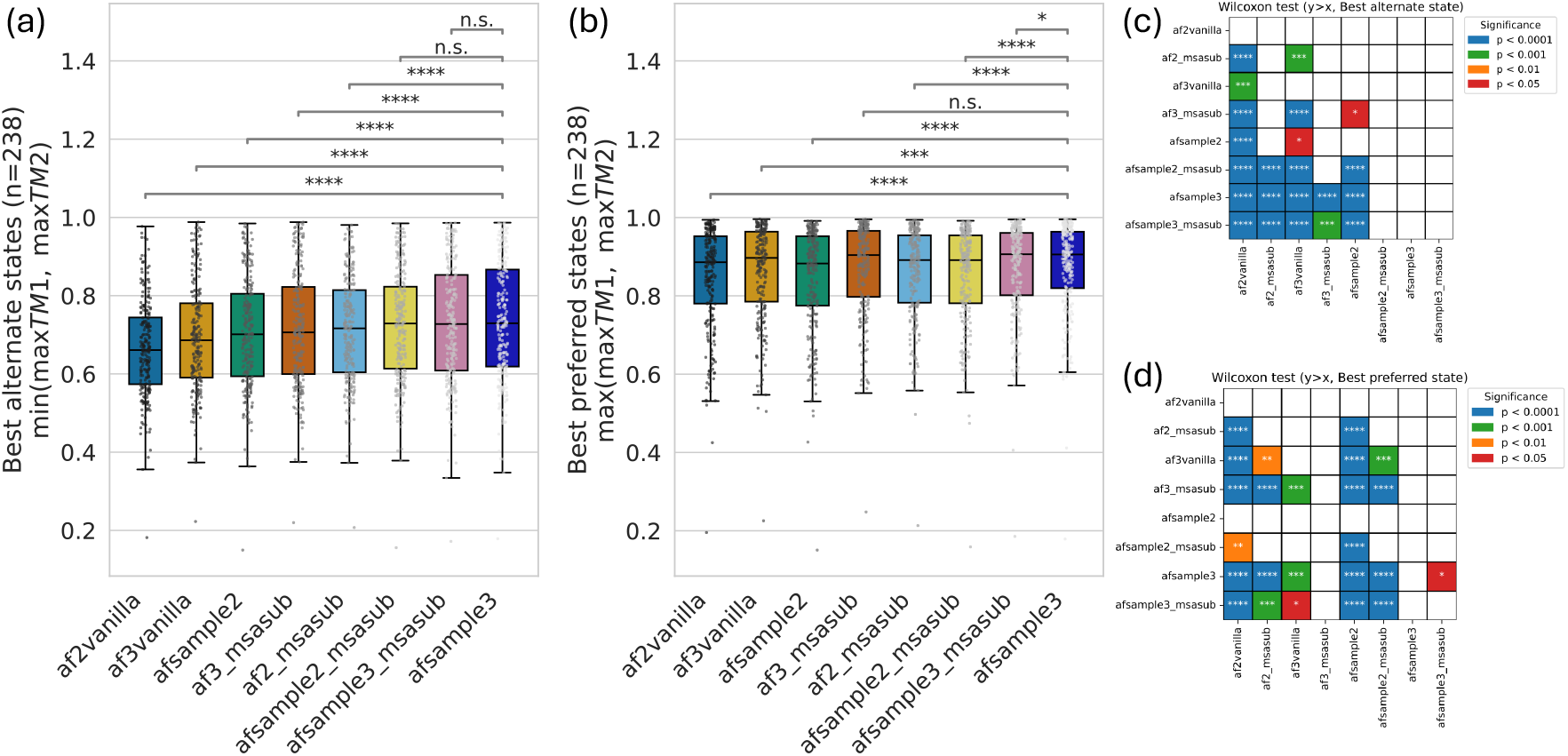
Comparing the effect of employing MSA subsampling on AF2vanilla, AF3vanilla, AFsample2, and AFsample3, for (a) best alternative state and (b) best preferred state, (c,d) pairwise Wilcoxon tests to ascertain if methods on the y-axis generate better states than those on the x-axis.

**Fig. 6.**
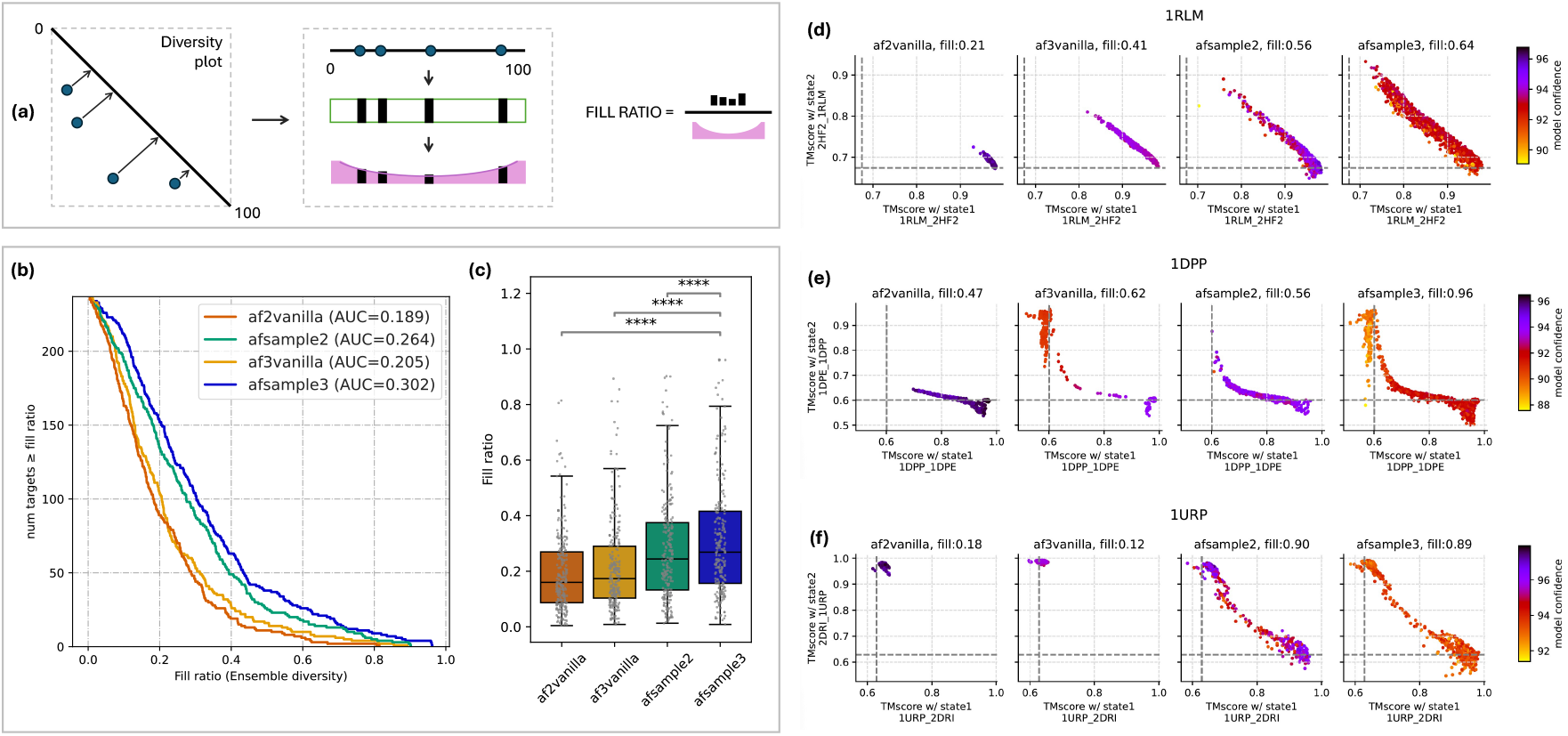
Quantifying and comparing ensemble diversity among methods. (a) Brief summary of the fill-ratio metric that aims at quantifying the diversity of a given protein ensemble while prioritizing the generation of end states. (b) Empirical cumulative distributions for all methods clearly demonstrate the improvements induced by AFsample2/3. (c) The overall distribution can also be visualized as a box-plot where AFsample3 is significantly batter than all other methods for generating diverse ensembles. (d-f) Diversity plots and the associated fill-ratios for three targets have been summarized. The overall change in the conformational signature is evident from both, the distribution of models in the ensemble, as well as the associated fill-ratio.

### Comparing ensemble diversity

The analysis thus far has primarily focused on the ability of the methods to generate distinct conformational states. However, evaluating ensemble characteristics, particularly structural diversity, is also important. To quantify ensemble diversity, we employed an enhanced version of the fill-ratio metric, originally developed for AFsample2 (5), that focus the attention on having end states before measuring the diversity.

While AF3vanilla showed improved state prediction accuracy relative to AF2vanilla, the corresponding gain in ensemble diversity was modest (mean *fill-ratio* - AF2vanilla: 0.195, AF3vanilla: 0.215; n=238). In contrast, the AFsample variants yielded substantially greater diversity, both in the inverse cumulative distribution profiles (Figure 6b,c) and in mean fill-ratios (AFsample2 = 0.279, AFsample3 = 0.313; n = 238; Table S1). It is important to note that all methods were evaluated using identical level of sampling, indicating that the observed differences in diversity stem primarily from the use of MSA perturbation rather than increased sampling. Table S1 summarizes the average fill-ratios for all methods across all 238 ensembles in the dataset.

To gain a more intuitive understanding of the generated ensembles, diversity plots for three representative protein systems (PDB IDs: 1RLM/2HF2, 1DPP/1DPE, and 1URP/2DRI) are shown in Figure 6d–f. These examples highlight that ensemble properties vary strongly depending on the method and modeling strategy.

For 1RLM, an E. coli sugar phosphatase SupH (Figure 6d), the baseline methods failed to recover either the alternate conformation (2HF2) or a broadly distributed ensemble. By contrast, AFsample2 and AFsample3 generated the alternate state and produced ensembles with substantially greater diversity.

For 1DPP, a periplasmic dipeptide transport/chemosensory receptor (Figure 6e), AF2vanilla did not recover the alternate state (1DPE), whereas AF3vanilla and AFsample3 both did. Nonetheless, AFsample3 generated a far richer ensemble (fill-ratio=0.96) compared to AF3vanilla (fill-ratio=0.62).

Finally, for 1URP, an E. coli ribose-binding protein (Figure 6f), neither AF2vanilla or AF3vanilla were even close to generating the alternate state, while both AFsample2 and AFsample3 successfully generated the alternate state, with relatively diverse ensembles.

Although only three illustrative cases are shown here, diversity plots for all 238 targets are provided in the Supplementary Information (Figure **??**). Overall, these results suggest that vanilla models are biased toward a dominant conformation, likely reflecting stronger evolutionary constraints encoded in the unperturbed MSA. Perturbation through masking (as in AFsample2 and AFsample3) appears to disrupt these constraints, enabling the model to escape from the preferred state and thereby generate more diverse ensembles. The potential physiological relevance of these putative intermediate states is further discussed in later sections.

### Investigating the validity of presumable intermediate states

The ability of the AFsample3 system to generate diverse protein ensembles was established in the previous sections. However, it is equally important to assess the relevance of the putative intermediate states within these ensembles. These intermediate conformations may represent transient but functionally critical states that facilitate biological processes such as ligand binding, allosteric regulation, or conformational transitions necessary for activity. Alternatively, they might simply be failed predictions of the stable end states, lacking biological significance. Distinguishing between these possibilities requires careful experimental validation and integration of complementary data to confirm whether these intermediates are genuinely accessed and functionally relevant in vivo. As an initial validation towards this end, the generated models were compared against experimentally determined structures in the Protein Data Bank (PDB), with the aim of identifying structural matches that serve as validation for true intermediate states in the structural ensembles.

Figure 7a provides an overview of the mapping of experimental PDB structures onto 118 diverse ensembles generated with fill-ratio>0.27. Out of these, 77 ensembles contained intermediate models that could be successfully mapped to known PDB structures, while 41 ensembles showed no structural correspondence. Figure 6b illustrates the latter case, where the ensembles span a wide range of conformational intermediates, yet only the end states coincide with experimentally determined structures. By contrast, for the ensembles depicted in Figure 6c, intermediate models could be mapped to the PDB, suggesting that at least some of the predicted intermediate conformations correspond to biologically relevant structural states.

**Fig. 7.**
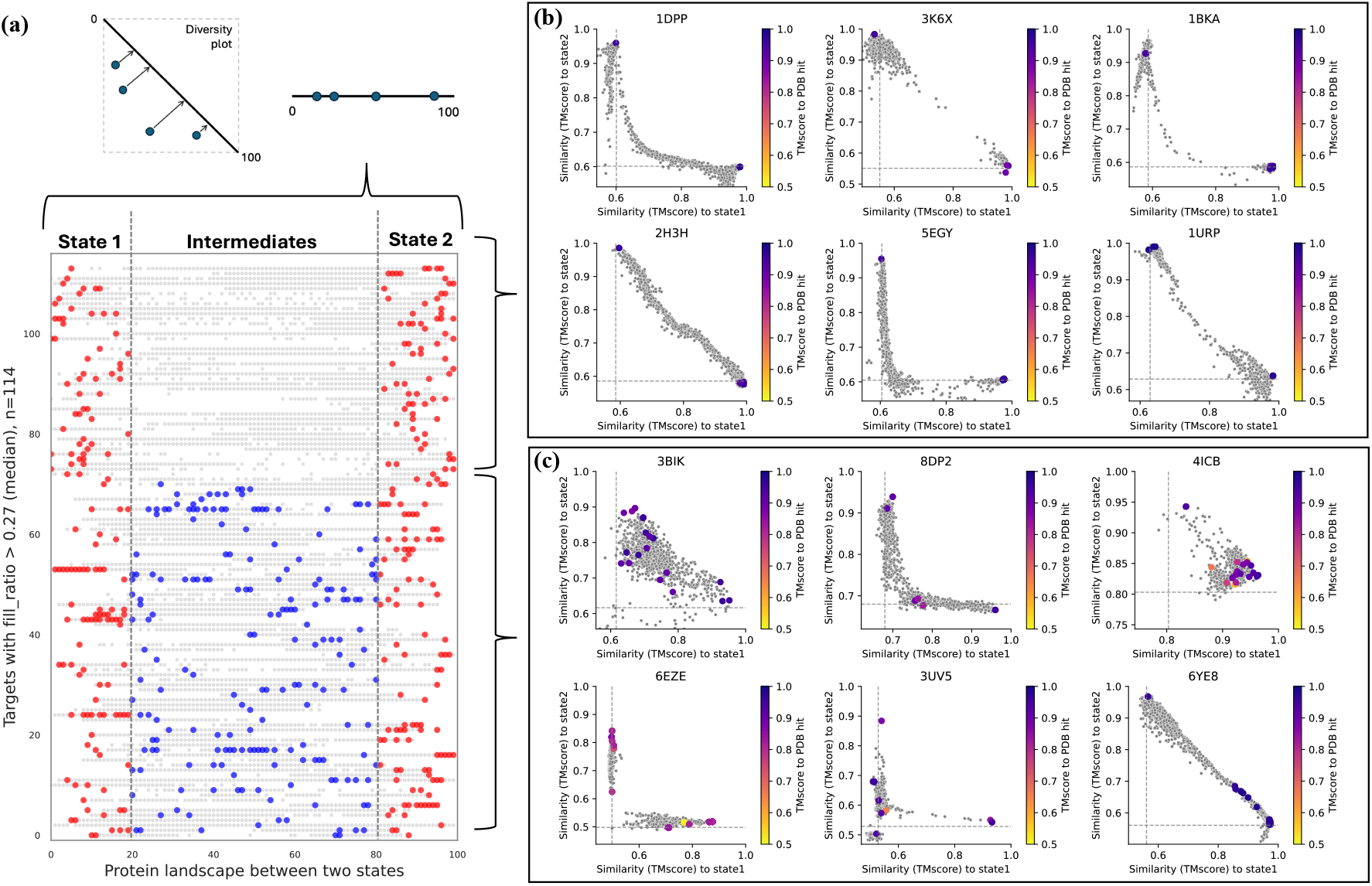
Investigating the validity of presumable intermediate states in AFsample3 ensembles. (a) Diversity plot flatten and visualized for all targets with a fill ratio > 0.27, gray points are models from the ensemble, and colored points are models mapped to the PDB, classified as close the either state (red) or intermediates (blue). (b) Examples of cases where no structure from the PDB could be mapped to the ensemble. (c) Examples of cases where the intermediates where mapped to the PDB.

Taken together, these mapped ensembles support the notion that MSA perturbations do not merely generate artificial variability, but rather enable the inference system to explore the natural conformational space of proteins. Importantly, the 41 ensembles for which no match could be identified should not be dismissed as irrelevant; instead, they may contain valid intermediate states that have not yet been experimentally solved. This further underlines the potential of such ensembles to extend beyond known PDB structures, offering mechanistic insights into functionally important, but so far uncharacterized, conformational transitions.

### Reference free state selection

All analysis up until now has been on the ability to generate structures similar to available experimental reference structures. In practical applications, however, such reference structures might not be available and it is therefore essential to establish a reliable strategy for identifying putative conformational states directly from predicted ensembles.

To identify distinct conformational states within each ensemble of predicted models, we extended the reference-free selection strategy introduced in AFsample2 (5) to provide a ranking of potential structural states. As outlined in Figure 8a, a pairwise TM-score similarity matrix was constructed for all models within each ensemble using Fold-seek (11). The resulting similarity matrix was subjected to principal component analysis (PCA) followed by *k*-means clustering to partition the ensemble into *k* structural clusters. The highest confidence model in a cluster was chosen as the cluster representative and ordered using a diversity-aware ranking scheme (DiSco): (1) the highest-confidence cluster was selected first; (2) the next representative was chosen as the cluster most dissimilar (i.e., lowest TM-score) to the previously selected one; and (3) the process was repeated iteratively until all clusters were ranked from 1 to *k*. Unless otherwise stated, *k* = 20 was used as the default number of clusters, as larger values did not affect performance.

**Fig. 8.**
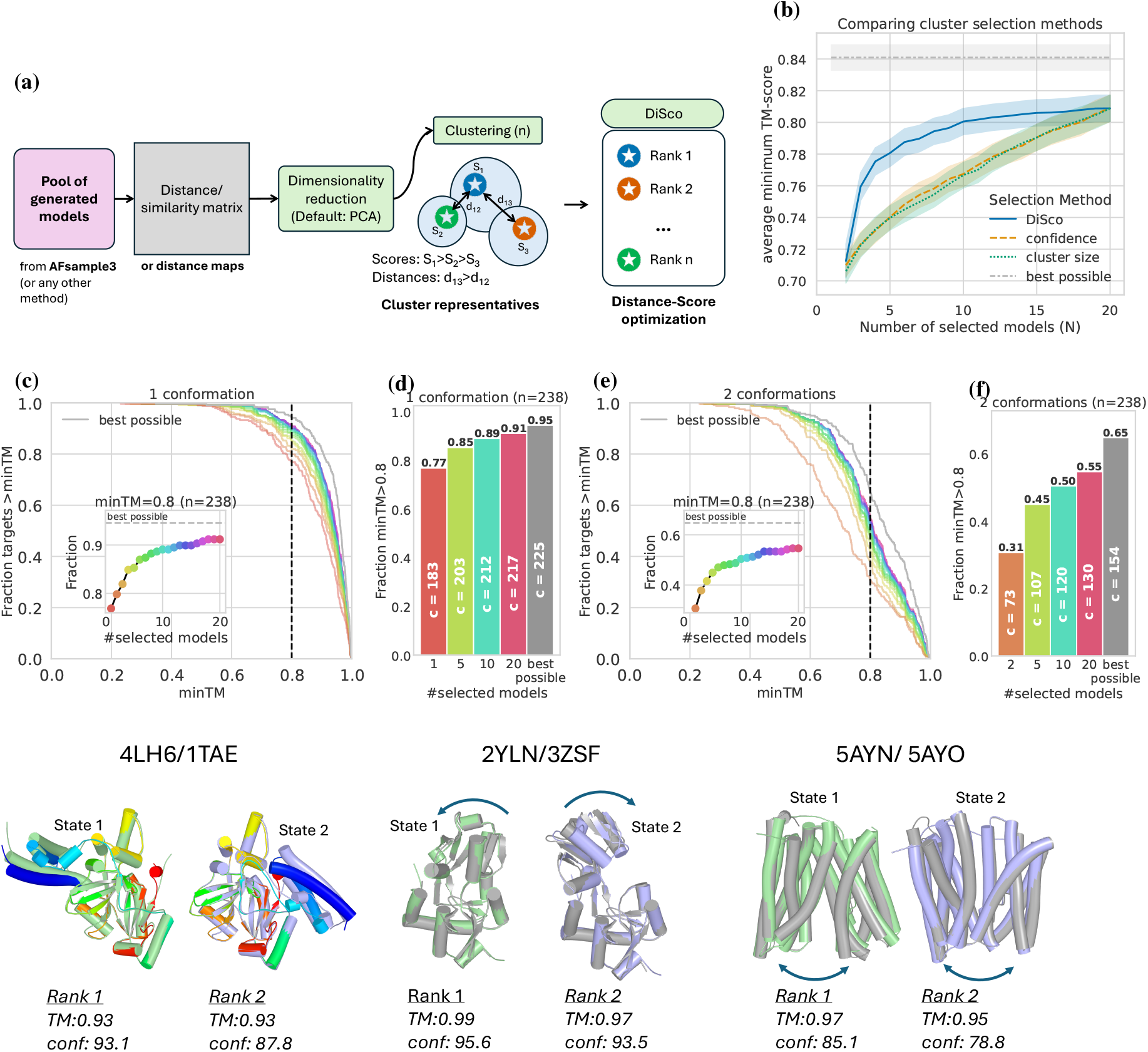
(a) Reference-free state selection protocol, given a protein ensemble, the aim is to identify possible conformational states. The system works on a similarity (or distance) matrix for all pairwise combinations of given models in the ensemble, clustering and distance-score optimization to generate a ranked list of representative models. (b) Comparing different methods of ranking the clusters from (a). (c) Inverse CDF for minTM when predicting 1 conformation when selecting rank 1-20 and best. (d) Fraction of targets with minTM>0.8 for different number of selected models when predicting 1 conformation. (e) Inverse CDF for minTM when predicting 2 conformations when selecting rank 1-20 and best. (f) Fraction of targets with minTM>0.8 for different number of selected models when predicting 2 conformations. (g) Selected successful examples with state 1 and state 2 (conformation 1 and conformation 2) predicted at rank 1 and rank 2.

Three different ways to order the *k* clusters were compared: DiSco, as described above, confidence from AlphaFold, and cluster size (Figure 8b). Clearly, DiSco selects signficantly better models compared to either using confidence or cluster size, which perform similar. At 5 selected models (top 5 cluster centers), the average minTM for the alternate state is 0.78 for DiSco and 0.74 for confidence and cluster size.

The ranked representatives were further assessed by analyzing the model quality, measured by minTM of the top 1 − *k* selected models when predicting a single state (Figure 8c,d), and for 2 − *k* selected models when predicting both states (Figure 8e,f). In single-state prediction (1 conformation), 77% (183/238) of the targets the a top-ranked model with TM>0.8. This proportion increases to 89% (212/238) when considering the top 10 ranked models, approaching the best possible selection of 95% (225/238). For predictions of both states, 33% of the targets have models with TM>0.8, which increases to 50% for top 10 models, relatively close to the best possible selection of 65% (154/238).

Together, these analyses demonstrate that the adapted reference-free selection scheme effectively identifies diverse conformational states within predicted ensembles while quantifying both complete and partial recovery performance across varying ensemble complexities

### Identifying potential novel conformations

The selection method is clearly ably to select good representative models in many cases. However, certain cases were observed where the top-ranked model from an ensemble did not resemble any of the known conformations for the target. Many of these “failed” selections with relatively high confidence could indeed be relevant structures that have not been experimentally characterized yet.

For example, in the case of the secreted frizzled-related protein (sFRP), which contains an N-terminal cysteine-rich domain (Fz domain) followed by a netrin-like domain and has two known conformations in the PDB (5XGP and 7EL5). AFsample3 successfully generated high quality models of 7EL5 with TM=0.88, but with lower confidence. The highest confidence model has a TM-score to any of the known conformation of around 0.55 (Figure 9a). Indeed the ensemble of all models seem to center around a TM-score of 0.55 to any of the states. The main difference between the models is the relative orientation of the Fz domain and netrin-like domain, resulting in an ensemble that samples the relative domain motion (Figure 9b,c). This observation aligns with the reported relative inter-domain orientation via hinge rotation between the cysteine-rich and netrin-like domains described in the original study and confirmed with SAXS experiments (12).

**Fig. 9.**
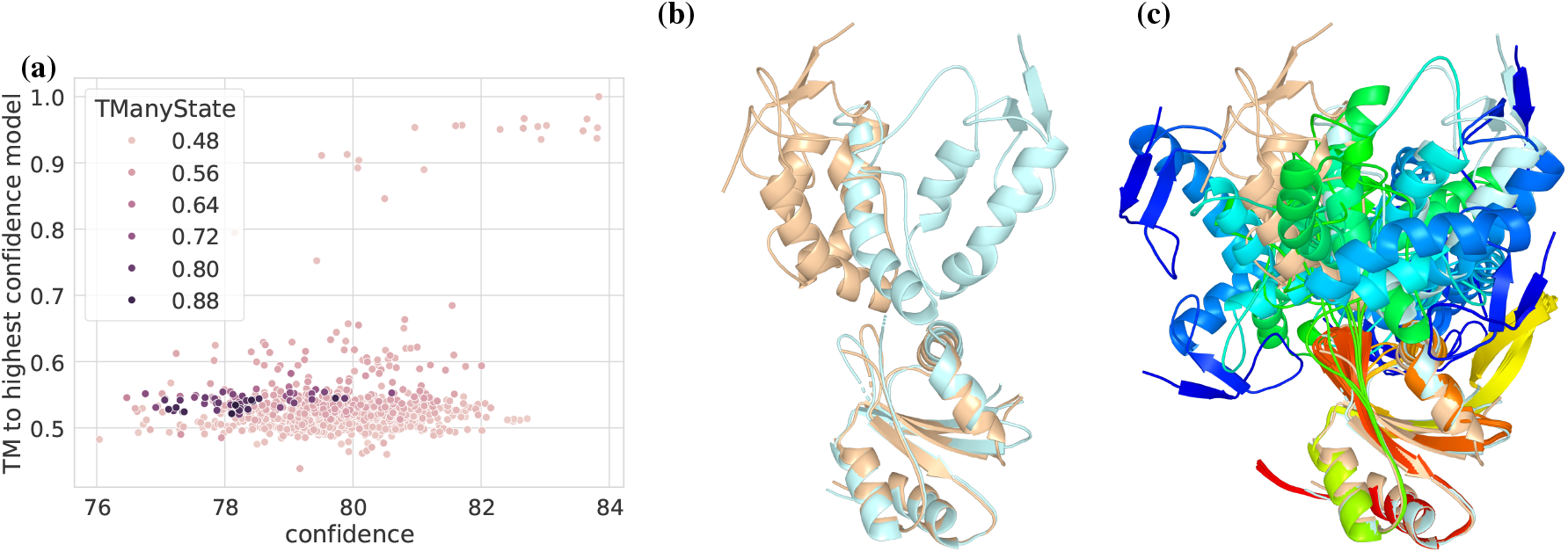
Example of a generated ensemble for the Szl protein (5XGP/7EL5). (a) Confidence vs TM-score to the highest confidence models, colored by the highest TM-score to any of the states (5XGP/7EL5). (b) 5XGP (wheat) and 7EL5 (palecyan) superimposed on the N-terminal domain. (c) The generated ensemble and native structures, superimposed on the N-terminal domain and colored in rainbow by chain.

## Discussion

Recent generative models for protein structure prediction have been very successful in generating accurate models for a diverse range of targets. However, predicting alternate state(s) remains a challenge. Methods such as MSA subsampling (2), AFsample (9), AFcluster (3), SPEACH_AF (4) and AFsample2 (5) have tried to solve the problem by essentially taking advantage of a very robust scoring function in Alphafold2. Most of these methods work by incorporating noise into the inference system in order to induce a conformational change and, with enough sampling, improve the chances of generating alternate states. Although existing methods have shown to work, the scope and application of the methods have always been limited by relatively small datasets.

Here, with AFsample3, we present a method to induce conformational diversity in generated models. It is based on the AF3 inference system and works by randomly masking columns of the MSA representation at runtime, with practically no computational overhead. We developed and evaluated AFsample3 on a large benchmark set of 238 targets, demonstrating substantial and statistically significant improvements in the generation of alternate conformations compared to existing methods. Additionally, we extend the reference-free state determination system to identify an arbitrary number of conformations from a given ensemble.

This study largely focuses on the ability of the methods in (i) generating alternate states, and (ii) generating diverse conformational ensembles. While the ability to generate accurate alternate states was quantified by comparing the best models with experimental structures from PDB. Ensemble diversity was evaluated with the in-house developed *fill-ratio* metric. In addition to this, a reference-free method to select presumable states from the generated protein ensemble was developed. This is particularly useful in the cases where experimental references are not available.

In addition, the functionality to run MSA randomizations in tandem with MSA subsampling was also added to AFsample2 and AFsample3. As shown in Figure S1, AFsample2 with MSA subsampling generated better alternate states than AFsample3 for 36 targets in the Cfold dataset. While AF-sample3 still performs better overall with 49 better targets, employing subsampling along with MSA masking was the optimal strategy for a sizable number of targets. Also, it should be noted that these improvements are not observed when MSA subsampling is employed in AF3 without any MSA masking, where AFsample3 is clearly better. This indicates that all benefits of MSA subsampling are already captured by MSA randomization for AF3.

It should be noted that while an ensemble of 1000 models was generated for each target, as observed in Figure 4b, the amount of sampling to generate the alternate state is much lower for majority of the targets. For most targets, generating 300 models was sufficient to get to the alternate state with AFsample3. This is a substantial improvement over AFsample2, where a similar level of performance is achieved at a higher sampling level. This points us to the conclusion that defining an optimal sampling level is still an unsolved problem. Even in the current set, an optimal sampling amount can range from as low as 10 models to as high as 750 models (Figure 4). One approach to solve this problem could be to track the diversity of the ensemble at runtime, which would add a computational overhead, but might indicate an optimal sampling level.

The definition of valid conformations in a protein system has always been as diverse as the conformations themselves. For instance, the initial dataset from Cfold had two conformations for every target in the set, however, the intermediate analysis identified multiple structures in the ensembles generated with AFsample3 that were successfully mapped to structures in the PDB.

While the performance of Alphafold3 (both vanilla and perturbed) is significantly better than Alphafold2 and its derivatives, it is very important to pin-point on the sources of these improvements. Improved model architecture, an optimized training strategy, amount of training data, or a combination of all might be a driver for the observed improvements. Also, recent studies have attributed the limited performance of AF2 on fold-switchers on memorization or overfitting (13). Similar studies might be required to test the limits of the AF3 inference system.

In summary, AFsample3 provides a significantly improved system to generate protein conformations. It was tested on a a large comprehensive set of proteins and performed considerable better than the previous version (AFsample2) in all metrics of performance relating to end state generation and ensemble diversity, while requiring less amount of sampling.

## Funding

This work was supported grants to BW from the Wallenberg AI, Autonomous System and Software Program (WASP) from Knut and Alice Wallenberg Foundation (KAW), Swedish Research Council grant, 2020-03352, 2024-05619. The Swedish e-Science Research Center. The computations were performed on resources provided by KAW and NSC (Berzelius).

## Supplementary Information

**Fig. S1.**
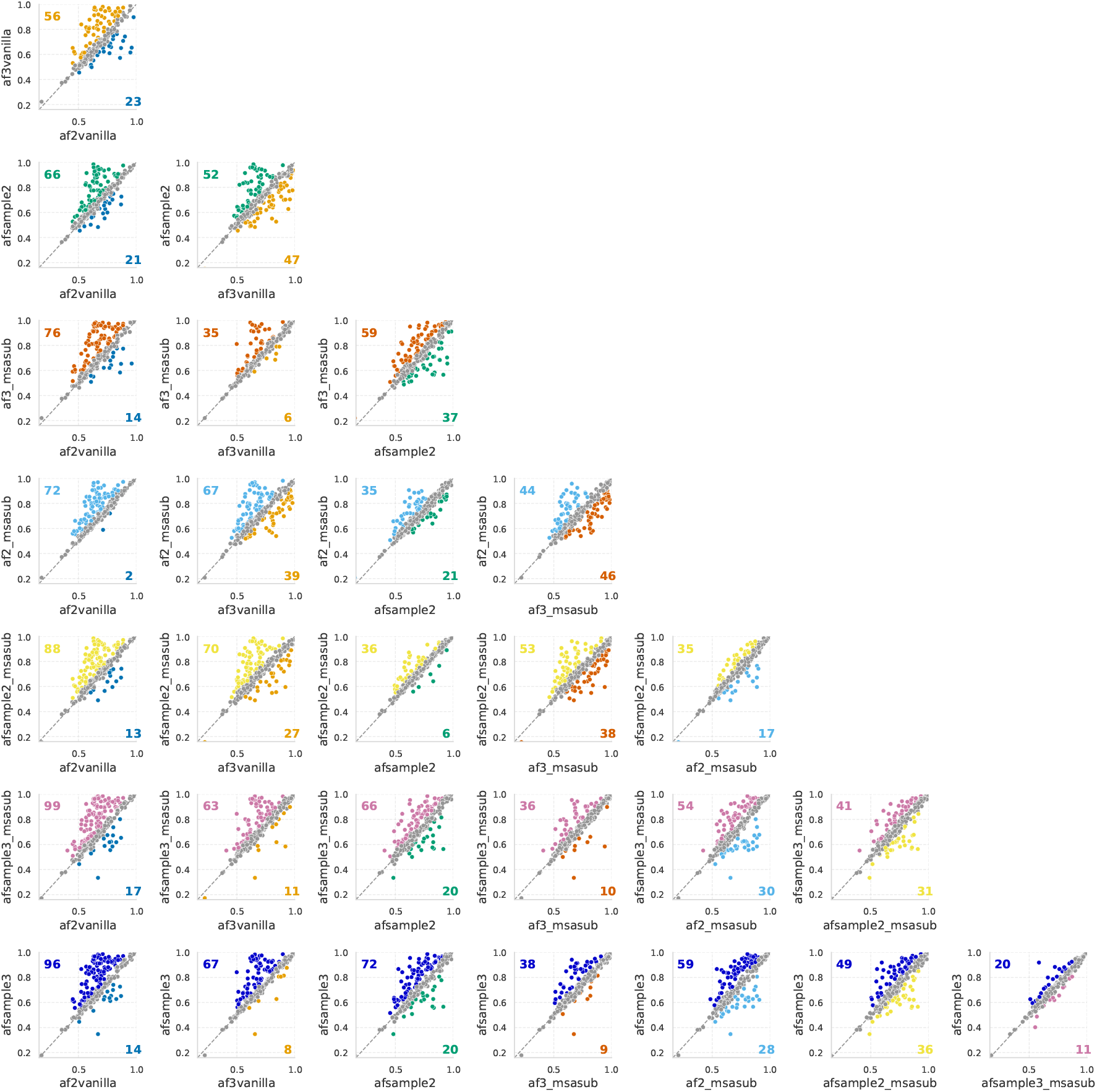
An all-v-all comparision of the best alternate states generated by all named methods under consideration. The definitions of the methods are summarized in Table S1

**Fig. S2.**
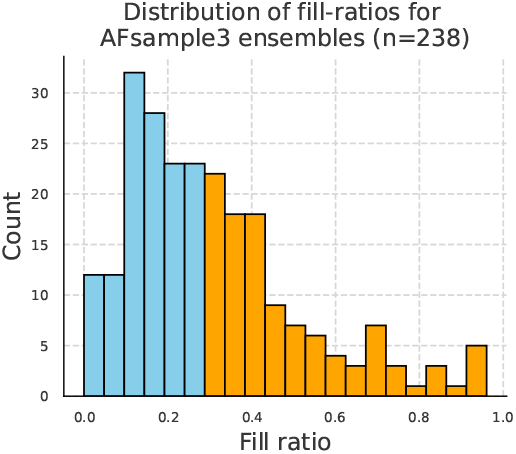
Distribution of fill ratios for all targets in the original cfold datasets for a two state system. Boxes are coloured based on if they are above or below the median (0.27).

**Table S1.** Experimental design and performance metrics for all AlphaFold variants. (*_m: shallow MSA implemented along with base method)

|  | af2vanilla | af3vanilla | afsample2 | afsample2_m | afsample3 | afsample3_m | af2_conformations | af3_m |
| --- | --- | --- | --- | --- | --- | --- | --- | --- |
| PARAMETERS |  |  |  |  |  |  |  |  |
| Neural net | AF2 | AF3 | AF2 | AF2 | AF3 | AF3 | AF2 | AF3 |
| Dropout | False | na | True | True | na | na | False | na |
| MSA randomization | 0% | 0% | 20% | 20% | 40% | 40% | 0% | 0% |
| MSA subsampling | False | False | False | True | False | True | True | True |
| PERFORMANCE METRICS |  |  |  |  |  |  |  |  |
| Fill ratio (ensemble diversity) | 0.195 | 0.215 | 0.279 | **0.339** | 0.313 | 0.323 | 0.307 | 0.262 |
| Mean TM (best alternate state) | 0.665 | 0.691 | 0.704 | 0.725 | **0.733** | 0.729 | 0.714 | 0.715 |
| Mean TM (best preferred state) | 0.847 | 0.862 | 0.843 | 0.854 | **0.874** | 0.870 | 0.860 | 0.869 |

## Notes

### Competing Interest Statement

The authors have declared no competing interest.

